# Analogues of the anti-malaria drug mefloquine have broad spectrum antifungal activity and are efficacious in a model of disseminated *Candida auris* infection

**DOI:** 10.1101/2024.08.29.610379

**Authors:** Soumitra Guin, Marhiah C. Montoya, Xiaoyu Wang, Robert Zarnowski, David R. Andes, Marvin J. Meyers, Noelle S. Williams, Damian J. Krysan

**Affiliations:** Department of Chemistry, School of Science and Engineering, Saint Louis University, Saint Louis, Missouri USA; Department of Pediatrics, Carver College of Medicine, University of Iowa, Iowa City, IA USA; Department of Biochemistry, UT Southwestern Medical Center, Dallas, Texas, USA; Department of Medicine, Section of Infectious Disease, University of Wisconsin, Madison, Wisconsin USA; Department of Medical Microbiology and Immunology, University of Wisconsin, Madison, Wisconsin USA; Department of Molecular Physiology and Biophysics, Carver College of Medicine, University of Iowa, Iowa City, IA USA

## Abstract

Only three classes of antifungal drugs are currently in clinical use. Here, we report that derivatives of the malarial drug mefloquine have broad spectrum antifungal activity including difficult to treat molds and endemic fungi. Pharmacokinetic and efficacy studies of NSC-4377 indicate it penetrates the central nervous system and is active against *Candida auris* in vivo. These data strongly support the further development of mefloquine analogs as a potentially new class of antifungal molecules.

---

The development of new antifungal drugs is a critical unmet clinical need (1). Currently, only three mechanistically distinct classes of antifungal drugs are available for the primary treatment of life-threatening fungal infections (2). The newest class of these medications, the echinocandins, was discovered in the 1970s and approved over twenty years ago. The urgent need for new mechanistic classes of antifungal drugs has been placed in stark relief by the emergence of pan-resistant strains of *Candida auris* (3). In addition to its high rates of resistance to both the azole and polyene classes of antifungal drugs, *C. auris* also has reduced susceptibility to widely used topical disinfectants such chlorhexidine (4). Resistance to these topical biocides is particularly important because *C. auris* persists on environmental surfaces and is transmissible from patient-to-patient as demonstrated by outbreaks in long-term care facilities.

The repurposing of medications approved to treat one condition as therapies for another condition has been widely investigated as a potentially expedient approach to identifying new drugs for difficult to treat diseases (5). Given the slow pace of new antifungal drug development, libraries of currently approved drugs have been extensively screened to identify candidates as either primary or adjunctive agents in the treatment of fungal infections (6). Many drugs have shown in vitro activity while a much smaller number have displayed efficacy in mammalian models of fungal infection. Furthermore, only two antifungal repurposing candidates, tamoxifen (7) and sertraline (8), have advanced to clinical trials. Both drugs were studied as adjuvants in combination with standard-of-care drugs for the treatment of cryptococcal meningitis (9, 10). Unfortunately, neither drug improved the efficacy of the standard therapy.

Although the direct use of a previously approved drug in the treatment of a new condition is the goal of most repurposing approaches, these candidates can play a second role in drug discovery. Specifically, the repurposed drug can serve as a hit with known human pharmacology upon which to base medicinal chemistry optimization of the new and desired activity. The latter approach to repurposing has not been widely applied to antifungal drug discovery. Our group’s work in this area has focused on derivatives of the breast cancer drug tamoxifen (7), phenothiazine antipsychotics (11) and the malaria drug mefloquine (12).

We reported previously that derivatives of the anti-malarial drug mefloquine generated at the Walter Reed Medical center as part of a project to optimize their utility for the treatment of malaria (Fig. 1A) have antifungal activity against human fungal pathogens including *C. auris* (12). Importantly, these molecules have activity against drug-resistant strains of *C. albicans, C. glabrata, C. auris*, and *Cryptococcus neoformans* suggesting that they are likely to have a novel mechanism of action. To further explore the spectrum of activity for this scaffold, the activity of NSC-2450 against a wide range of human fungal pathogens was characterized through the NIH Contract testing lab. As shown in Table 1, NSC-2450 showed a wide spectrum of activity including the previously mentioned yeasts, difficult to treat molds (*Rhizopus, Fusarium, Scedosporium*) and endemic fungi including *Coccidioides*. Although the minimum inhibitory concentration (MIC) for NSC-2450 were modest against some of these organisms (16-32 µg/m), the data establish the broad spectrum of activity of the general scaffold.

**Figure 1.**
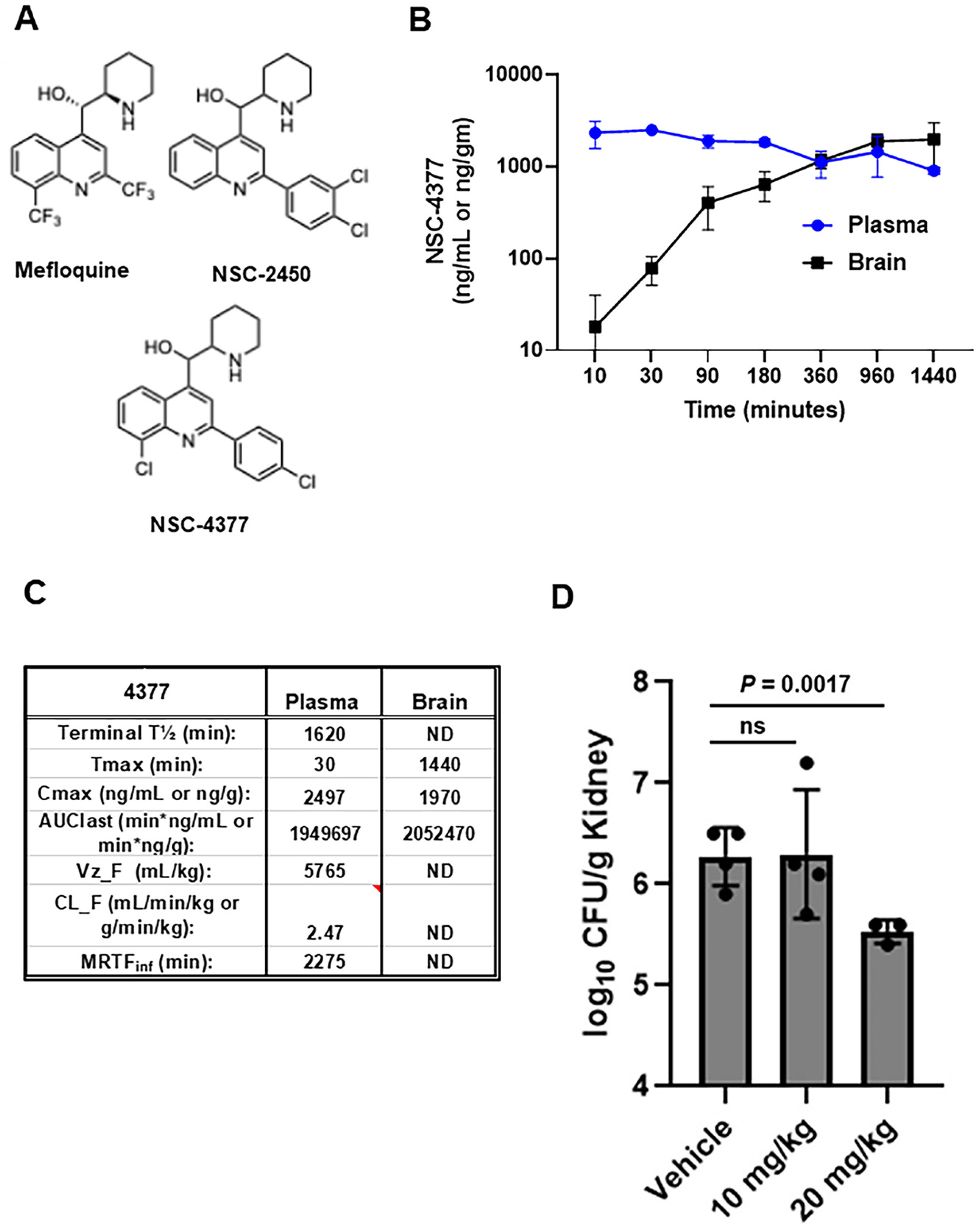
Pharmacokinetics and efficacy of NSC-4377 against *Candida auris* in mouse models. **A**. Structures of mefloquine, NSC-2450, and NSC-4377. **B**. Plasma and brain concentrations of NSC-4377 in mice. Points are mean of three mice with standard deviation shown by error bars. **C**. Pharmacokinetic parameters for NSC-4377 derived from data shown in (**B**). **D**. Fungal burden of kidneys 72 hr post-infection for neutropenic mice treated daily with vehicle, 10 mg/kg, or 20 mg/kg NSC-4377. Data are mean and standard deviation for 3-4 mice per group.

**Table 1.**
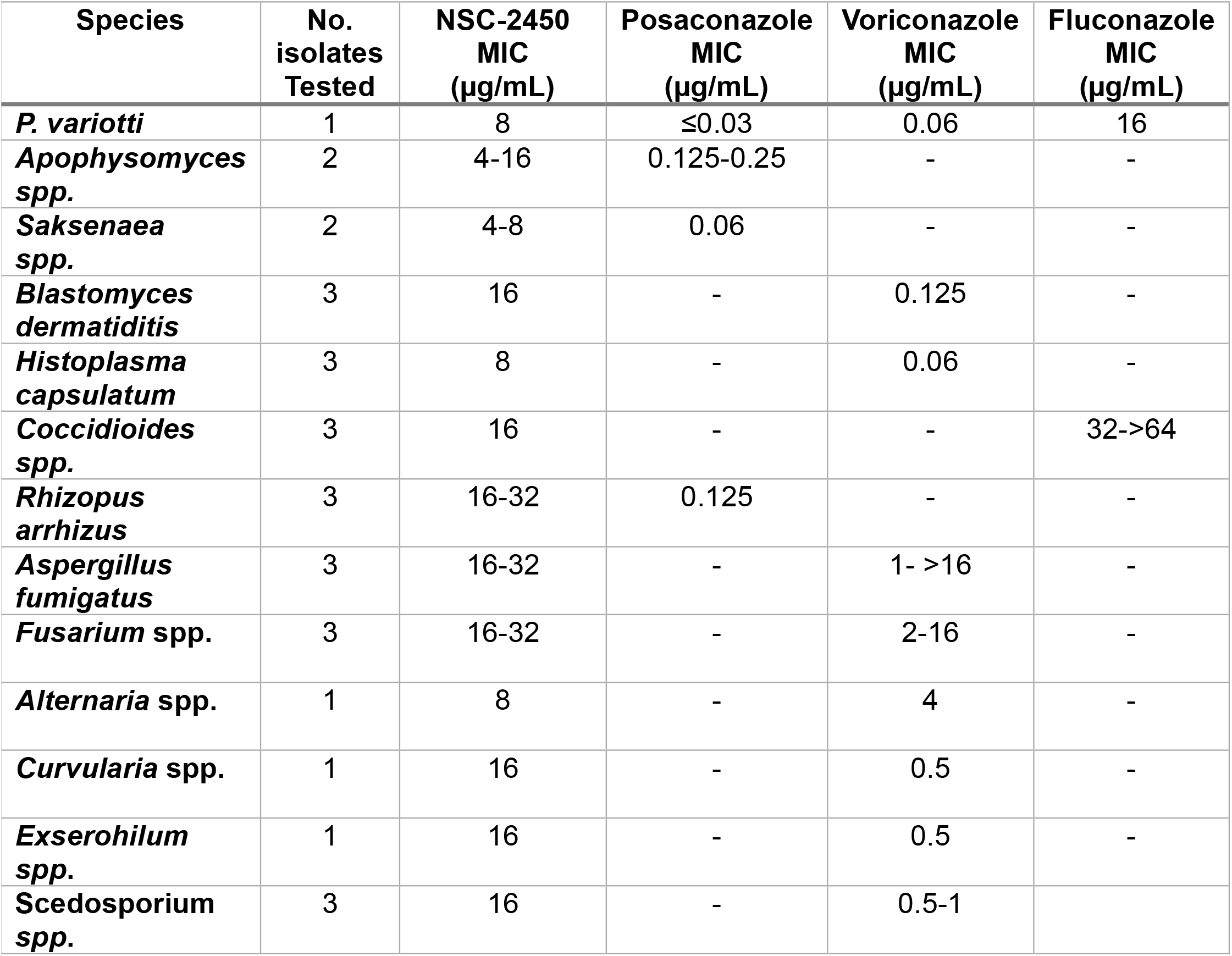
Spectrum of antifungal activity for NSC-2450.

| **Species** | **No. isolates Tested** | **NSC-2450 MIC (µg/mL)** | **Posaconazole MIC (µg/mL)** | **Voriconazole MIC (µg/mL)** | **Fluconazole MIC (µg/mL)** |
| --- | --- | --- | --- | --- | --- |
| *P. variotti* | 1 | 8 | ≤0.03 | 0.06 | 16 |
| *Apophysomyces spp.* | 2 | 4-16 | 0.125-0.25 | - | - |
| *Saksenaea spp.* | 2 | 4-8 | 0.06 | - | - |
| *Blastomyces dermatitidis* | 3 | 16 | - | 0.125 | - |
| *Histoplasma capsulatum* | 3 | 8 | - | 0.06 | - |
| *Coccidioides spp.* | 3 | 16 | - | - | 32->64 |
| *Rhizopus arrhizus* | 3 | 16-32 | 0.125 | - | - |
| *Aspergillus fumigatus* | 3 | 16-32 | - | 1- >16 | - |
| *Fusarium spp.* | 3 | 16-32 | - | 2-16 | - |
| *Alternaria spp.* | 1 | 8 | - | 4 | - |
| *Curvularia spp.* | 1 | 16 | - | 0.5 | - |
| *Exserohilum spp.* | 1 | 16 | - | 0.5 | - |
| *Scedosporium spp.* | 3 | 16 | - | 0.5-1 |  |

Next, we took advantage of an efficient synthetic route to NSC-4377 recently developed by our group (13) to characterize its pharmacokinetic/pharmacodynamic (PK/PD) properties as well as it’s efficacy in a mouse model of disseminated *C. auris*. We first characterized the serum and brain PK/PD parameters of NSC-4377 in CD-1 mice (3 mice per time point) after a 10 mg/kg intraperitoneal (IP) dose (Fig. 1B&C) by LC/MS-MS as described previously (14). Overall, the plasma and brain exposures of NSC-4377 were comparable with C_*max*_ of ∼2 µg/mL(g) in both compartments (Fig. 1C). These data are similar to the plasma and brain PK/PD characteristics reported by IV administration of a 5 mg/kg to FVB mice reported by Dow et al (15). A single 10 mg/kg dose provides plasma concentrations near the MIC of NSC-4377 yeast such as *C. albicans, C. auris*, and *C. neoformans*. Since the brain is a target organ for multiple fungal pathogens and the primary target organ for *C. neoformans*, which causes 150,000 deaths/year (16), the high concentrations in the brain are important feature of this class of molecules. In addition, the long half-life of NSC-4377 suggests that multiple doses are likely to establish drug concentrations above MIC for a broad spectrum of fungal pathogens.

Based on the promising PK/PD characteristics for NSC-4377, we tested its efficacy in a neutropenic mouse model of disseminated *C. auris* infection. As previously described (17), we used the clade IV *C. auris* strain B11804. Mice were infected, treated with two doses (10 or 20 mg/kg) of NSC-4377 and kidneys were harvested 72 hr post-infection and the fungal burden was determined by quantitative plating on YPD plates incubated at 30°C for 24hr. Although the 10 mg/kg dose did not have an effect on kidney fungal burden, treatment with 20 mg/kg led to a statistically significant reduction in the kidney fungal burden (Fig. 1D). The 10 and 20 mg/kg doses were well-tolerated while dosing at 40 mg/kg led to signs of toxicity.

Taken together, the broad spectrum of antifungal activity, the PK characteristics and in vivo efficacy data reported herein indicate that the mefloquine-derived, amino-quinoline scaffold has attractive features that support further pre-clinical optimization and development. Indeed, NSC-4377 is now one of the few novel antifungal molecules that display in vivo activity against *C. auris*, a fungal pathogen which can have no therapeutic options.

## Acknowledgements

This was supported in part by R21AI164578 (MJM&DJK). We thank Jeniel Nett (Wisconsin) for additional experiments.

